# Hepatic lipopolysaccharide binding protein partially uncouples inflammation from fibrosis in MAFLD

**DOI:** 10.1101/2024.06.17.599212

**Authors:** Dan Wang, Ania Baghoomian, Zhengyi Zhang, Ya Cui, Emily C. Whang, Xiang Li, Josue Fraga, Rachel Ariana Spellman, Tien S. Dong, Wei Li, Arpana Gupta, Jihane N. Benhammou, Tamer Sallam

**Affiliations:** Division of Cardiology, Department of Medicine, University of California, Los Angeles, CA; Department of Physiology, University of California, Los Angeles, CA; Molecular Biology Institute, University of California, Los Angeles, CA; Division of Computational Biomedicine, Biological Chemistry, University of California, Irvine, CA; Department of Pathology and Laboratory Medicine, University of California, Los Angeles, CA; Department of Biological Chemistry, University of California, Los Angeles, CA; Pediatric Gastroenterology, Hepatology, and Nutrition, David Geffen School of Medicine, University of California, Los Angeles, CA; Vatche and Tamar Manoukian Division of Digestive Diseases, University of California, Los Angeles, CA; Division of Gastroenterology, Hepatology and Parenteral Nutrition, Department of Medicine, VA Greater Los Angeles Healthcare System, CA; Comprehensive Liver Research Center, University of California, Los Angeles, CA; Goodman-Luskin Microbiome Center, University of California, Los Angeles, CA; G. Oppenheimer Center for Neurobiology of Stress and Resilience, University of California, Los Angeles, CA; Jonsson Comprehensive Cancer Center, University of California, Los Angeles, CA

## Abstract

Non-alcoholic fatty liver disease (NAFLD), recently renamed metabolic-associated fatty liver disease (MAFLD), is the most common liver disease worldwide. The progression to fibrosis, occurring against a backdrop of hepatic steatosis and inflammation, critically determines liver-related morbidity and mortality. Inflammatory processes contribute to various stages of MAFLD and thought to instigate hepatic fibrosis. For this reason, targeting inflammation has been heavily nominated as a strategy to mitigate liver fibrosis. Lipopolysaccharide binding protein (LBP) is a secreted protein that plays an established role in innate immune responses. Here, using adoptive transfer studies and tissue-specific deletion models we show that hepatocytes are the dominant contributors to circulating LBP. In a murine model of MAFLD, hepatocyte-specific deletion of LBP restrained hepatic inflammation and improved liver function abnormalities, but not measures of fibrosis. Human studies, including genetic evidence, corroborate an important role for LBP in hepatic inflammation with minimal impact on fibrosis. Collectively, our data argues against the idea that targeting hepatic inflammation is a viable approach to reducing fibrosis.

---

Metabolic dysfunction-associated fatty liver disease (MAFLD) is a heterogeneous spectrum liver disorder affecting 20% of the population. Metabolic dysfunction-associated steatohepatitis (MASH) is a more advanced form associated with inflammation and fibrosis, which can progress to cirrhosis. Scarring of liver tissue is a strong predictor of poor clinical outcomes and multiple factors act synergistically to license fibrosis. Of particular interest is the role of inflammation, since the “multiple hit” model of MASH implies that inflammation incites fibrosis (1). Thus, targeting inflammation has been proposed to combat hepatic fibrosis. Surprisingly, treatment with the C-C chemokine receptors (CCR) antagonist, cenicriviroc, failed to meaningfully change fibrosis in a recent Phase III clinical trial (2). These results and others have raised questions about the therapeutic potential of targeting inflammation in MASH and our understanding of sequential events leading to the progression of MASH.

To investigate the interplay between innate immune responses and fibrosis, we explored a role for LBP in MAFLD. LBP is known to be induced in response to inflammatory signaling and facilitates immune cell recruitment and function(3, 4). Feeding mice diets known to induce MAFLD or treatment with LPS increased circulating LBP (Figure 1A, Supplemental Figure 1, A-D). LBP is expressed in different tissues, but the main source of circulating LBP levels remains unknown. LBP expression is highest in the liver and specifically parenchymal cells (Supplemental Figure 1, E-F), which led us to the hypothesize that hepatocytes are the key contributors to circulating LBP. To explore this, we generated LBP^flox/flox^ mice (Supplemental Figure 1G) and observed that hepatocyte-specific loss of LBP using an AAV-TBG-Cre approach completely abolished circulating LBP levels (Figure 1B). In addition, immune reconstitution of wildtype (WT) or LBP^-/-^ bone-marrow (4) on a hyperlipidemic background, did not show changes in circulating LBP levels (Figure 1C). Collectively, these results suggest that circulating LBP is predominantly dictated by hepatocytes.

**Figure 1.**
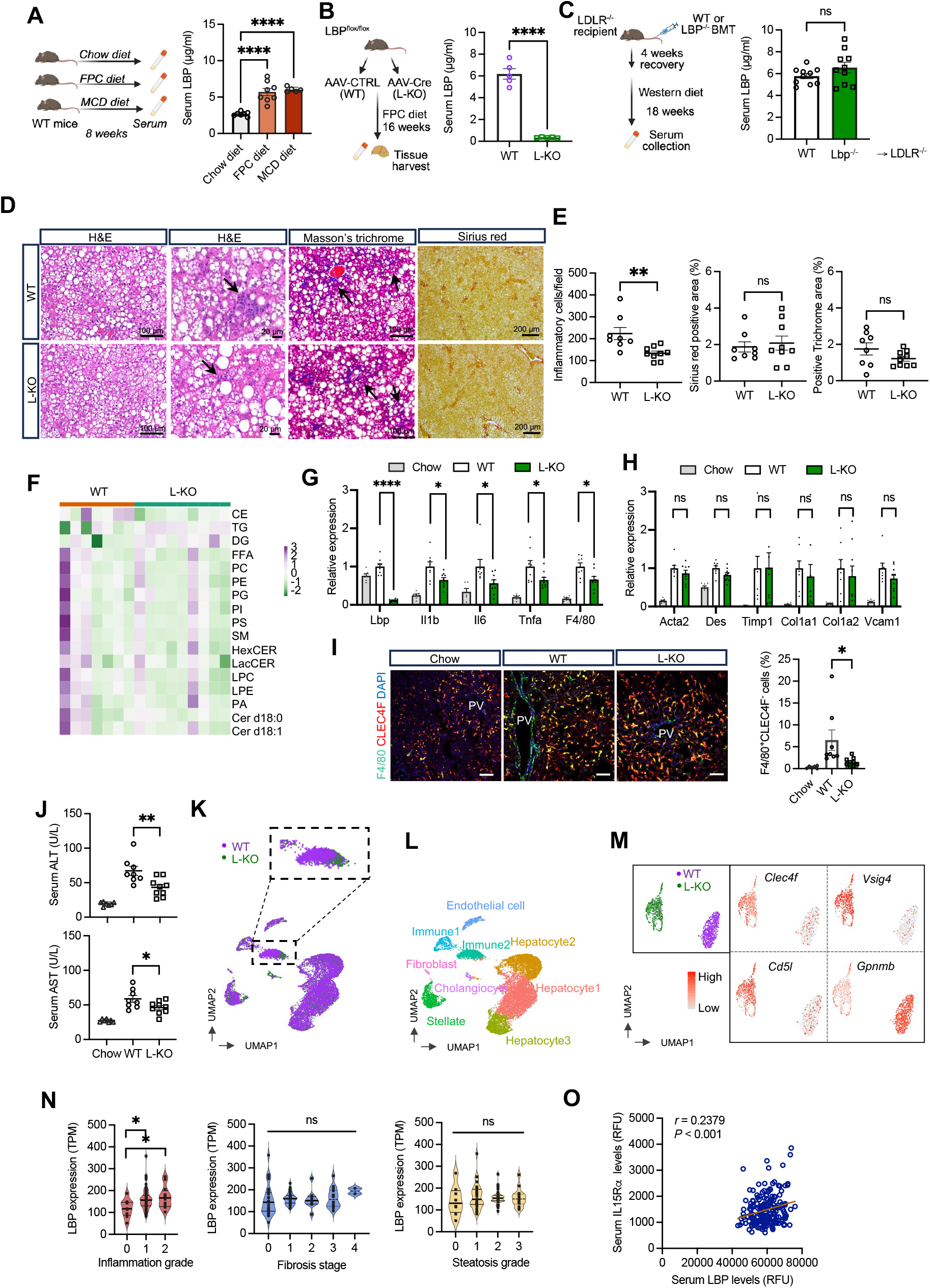
Hepatic LBP deficiency reduces inflammation but not fibrosis. (**A**-**C**). Serum LBP in male mice as indicated. (n= 5-8 in (A), n= 5 in (B) and n= 10 in (C)). (**D**). Representative liver H&E staining (arrows indicate inflammatory cell infiltration), Masson’s trichrome staining and Sirius red staining images. (**E**). Quantification of inflammatory cells and fibrosis. (**F**). Liver lipidomics heatmap. (**G**-**H**). RT-qPCR from liver. (**I**). Liver immunohistochemistry, Bar 100 μm. (**J**). Serum ALT and AST. (n=8-9 in (E-J)). (**K**). UMAP plot of snRNA-seq from liver. (**L**). Annotation of populations derived from cluster-specific gene expression analysis. (**M**). Representative genes in UMAP plot from Immune2. (**N**). LBP expressions in human liver. (**O**). Correlations between circulating LBP and serum IL15Ra in human MASH cohort. Data represent mean ± SEM. Non-paired Student’s *t*-test. *: p < 0.05; **: p < 0.01; ****: p < 0.0001; ns: not significant.

To better understand the role of hepatocyte LBP in MAFLD, we fed hepatocyte-specific LBP knockout mice (L-KO) or controls (WT) FPC (rich in fructose, palmitate, and cholesterol) diet. We did not observe differences in hepatocyte lipid droplet, lipid species composition (Figure 1D-F), animal weight, percent fat, and liver weight between groups (Supplemental Figure 2, A-D). LBP deficiency led to a significant reduction in inflammatory cells (Figure 1, D-E) compared with controls, although we did not observe differences in fibrosis as assessed and quantified by Masson-trichrome, Sirius red and αSMA staining (Figure 1, D-E, Supplemental Figure 2E). In line with the above results, gene expression analysis showed a reduction in inflammatory markers (Figure 1G) in L-KO compared to controls without changes in fibrosis (Figure 1H, Supplemental Figure 2F) or lipid metabolism genes (Supplemental Figure 2, G-H). Furthermore, infiltrating monocytes/macrophages were reduced in L-KO compared to controls as shown by F4/80^+^CLEC4F^-^ (Figure 1I) and Ly6C^+^ staining (Supplemental Figure 2E), along with a significant reduction in circulating and liver inflammatory markers (Supplemental Figure 2, I-J). Transaminases were significantly elevated in controls compared with L-KO (Figure 1J), suggesting that the observed change in inflammation meaningfully impacts liver steatohepatitis. Taken together, the results suggest that loss of LBP reduces inflammatory activation in MAFLD independent of hepatic lipid composition and without altering fibrosis.

To confirm the influence of LBP deficiency on hepatic immune cell composition, we performed single-nucleus RNA-Seq (snRNA-Seq) on livers from WT and L-KO mice. Integrated transcriptomic analysis revealed distinct population liver cell-type clusters (Figure 1, K-L, Supplemental Figure 2K). No differences were seen in the expression of key fibrosis genes in stellate cells (Supplemental Figure 2L). Immune1 cluster included kupffer cells, neutrophils, and dendritic cells, as suggested by high cell type-specific markers including *Adrge1* (Supplemental Figure 2K). Major changes between WT and L-KO centered in the Immune2 cluster (Figure 1, K-L). This cluster was enriched in recruited macrophage populations expressing low/intermediate macrophage markers including *Adrge1* (F4/80) and *Csf1r* (Supplemental Figure 2K). Further analysis of this cluster revealed remarkably distinct populations (Figure 1M, upper left). L-KO subcluster showed enrichments of non-inflammatory macrophage markers including *Vsig4*, *Cd5l* and *Clec4f* (Figure 1M) and anti-inflammation genes like *Lrg1* and *Gna15* (Supplemental Figure 2M), whereas WT exhibited higher levels of *Gpnmb*, which defines pro-inflammatory macrophages (Figure 1M).

To confirm the impact of LBP on hepatic scarring, we treated WT or L-KO mice with carbon tetrachloride (CCl_4_) and did not observe differences in fibrosis (Supplemental Figure 3A-E). To translate our findings to human MAFLD, we first explored the expression of human LBP from GTEx. Human LBP is dominantly expressed in the liver and specifically hepatocytes (Supplemental Figure 3, F-G). In a cohort of MASH patients, we found that hepatic LBP expression strongly segregates the degree of liver inflammation but not steatosis or fibrosis (Figure 1N). GWAS showed that coding-mutations in LBP are strongly associated with circulating markers of inflammation, known to be important in MAFLD-like serum IL-15 levels (*P* value 8 x 10^-163^) (Supplemental Figure 3H). We confirmed a positive correlation between circulating IL-15Rα and LBP levels in an independent cohort of MASH patients (Figure 1O). Taken together, our results suggest that the human LBP may be protective against hepatic inflammation with minimal impact on fibrosis.

In summary, we uncover a previously unidentified role for hepatic LBP in MAFLD. We find that loss of LBP reduced inflammation along with macrophage recruitment markers, but these changes are not sufficient to reduce hepatic fibrosis. Since hepatic scarring is a crucial driver of liver disease-related morbidity and mortality, our findings have important implications for approaches that aim to target inflammation to reduce fibrosis.

## Supplemental Material

### Methods

#### Animals and diets

All mice used in this study were housed in pathogen-free rooms with temperature monitors and a 12-hour light/12-hour dark cycle. LBP mice were generated from the EUCOMM/KOMP-CSD ‘Knockout-First’ C57BL/6 ES cell resource and then crossed with the FLPe knock-in mouse strain (JAX 009086) to generate the LBP^f/f^ mice. Hepatocyte-specific LBP knockout mice were injected with AAV8-TBG-Cre (L-KO) or AAV8-TBG-Control (WT) (Addgene 107787). The diets used were the standard chow diet (Research Diets), FPC-MASH diet (Envigo, TD.190142), Western diet (Research Diets, D12079B), MCD diet (Research Diets, A02082002B), and mice were fed as indicated. The bone marrow transplantation was performed as previously described (4). Intraperitoneal injection of lipopolysaccharide (LPS) at 1.5mg/kg for a single dose, or daily injection for up to 7 days. For CCl_4_-induced liver fibrosis, L-KO and WT mice were injected i.p. biweekly for 4 weeks with 0.5 μl/g body weight of CCl_4_ (Sigma), which was dissolved in corn oil at a ratio of 1:3. We did not observe differences between males and females in pilot studies and male mice were used in presented data.

#### Histopathological Analysis and Immunofluorescence

Hematoxylin and eosin (H&E) staining and Masson’s trichrome staining of liver sections were prepared and stained from paraffin-embedded sections and performed by the Translational Pathology Core Laboratory (TPCL) at UCLA. For Sirius red staining, the embedded samples were cut into 4 μm-thick sections and stained with Sirius Red (saturated picric acid containing 0.1% DirectRed 80, Sigma-Aldrich). For assessing the fibrosis in Masson’s trichrome staining and Sirus red staining, the sections were evaluated using x5 magnification. 4∼5 different microscopic fields were selected randomly in each individual liver section. The inflammatory cells in H&E staining were quantified as the number of mononuclear cells per field using a 20x objective. For each slide, 3∼5 different microscopic fields were selected randomly in each individual liver section. The inflammatory cells per field and fibrosis contents from each mouse were quantified with ImageJ software. For immunofluorescence, liver sections were labeled with primary antibodies overnight at 4 °C, followed by incubation with a fluorophore-conjugated secondary antibody for 1 hour. Primary antibodies are anti-F4/80 at a 1:400 dilution, Abcam, ab6640, clone CI:A3-1; anti-alpha-smooth muscle actin monoclonal antibody at a 1:400 dilution, eBioscience, clone 1A4; anti-Ly-6C at a 1:200 dilution, BD Pharmingen, Clone: HK1.4; and anti-CLEC4F/CLECSF13 antibody at a 1:200 dilution, R&D. The stained sections were stained with DAPI and then mounted (Life Technologies, P36935).

#### Serum LBP ELISA

Blood samples were collected and kept cool on ice. The serum was collected after centrifuge for 10 minutes at 8,000g at 4 °C. The samples were stored at −80 °C. Before conducting the assay, the samples were brought to room temperature and gently mixed to avoid foaming. The LBP ELISA was performed according to the manufacturer’s instructions (HycultBiotech, HK205). Briefly, 100 μl of the diluted sample and standards were transferred to the appropriate wells. The plate was incubated at room temperature for one hour before being washed with 200 μl of dilution buffer. Then 100 μl of the tracer and 100 μl of the streptavidin-peroxidase were added in sequence and incubated for 1 hour in each reaction. After washing, the TMB substrate was incubated for 20 minutes before adding stop solutions. The plate reader was set at 450 nm following the instructions, and the plates were read with readings performed within 30 minutes of adding the stop solution.

#### Liver lipidomics analysis

For lipid quantification analysis, approximately 80 mg of frozen liver was used. Samples were analyzed in UCLA Lipidomics Core on the Sciex Lipidyzer Platform using mass spectrometry methodology for targeted quantitative measurement of over 1400 lipid species across 17 lipid subclasses (5, 6). The lipid subclasses include cholesterol esters (CE), cerimides (Cer d18:1), diacylglycerol (DG), dihydroceramides (Cer d18:0), free fatty acids (FFA), hexosylceramides (HexCER), lactosylceramide (LacCER), lysophosphatidylcholine (LPC), lysophosphatidylethanolamine (LPE), phosphatidic acid (PA), phosphatidylcholine (PC), phosphatidylethanolamine (PE), phosphatidylglycerol (PG), phosphatidylinositol (PI), phosphatidylserine (PS), sphingomyelin (SM), and triacylglyceride (TG). Each quantitative value was normalized to milligrams of corresponding samples.

#### Gene expression analysis (RT-qPCR)

The liver samples were homogenized and prepared for RNA isolation using TRIzol (Invitrogen) (7). The cDNA was synthesized using reverse transcription enzyme with oligo dT and random hexamers. Quantitative real-time PCR (qRT-PCR) was performed using SsoAdvanced Univ SYBR Green (Bio-Rad 1725275) to analyze gene expression. The results were normalized to housekeeping gene 36B4. The primer sequences for qRT-PCR are listed in Supplemental Table 1.

#### Immunoblotting

About 100 mg of the protein from the frozen liver tissue samples was isolated and extracted. The samples were dounced using 300 μl of RIPA buffer (Boston Bioproducts) and centrifuged for 15 minutes at 10,000 g. The pure protein layer was extracted without disturbing the pellet or fat layers. Then, the protein concentration was quantified utilizing BCA assay. Equal amounts of protein were loaded into the 4-12% NuPAGE Bis-Tris Gel (Invitrogen) and the gel was run at 130 volts for about 1 hour and 30 minutes. The protein was transferred from the gel to the PVDF membrane (0.45μm) at 100 volts and then blocked in a 5% milk solution. The primary antibodies were anti-Phospho-p44/42 MAPK (Erk1/2) (Thr202/Tyr204) Antibody (Cell Signaling Technology, 9101) and anti-GAPDH (GeneTex, GTX627408). The protein quantification was performed using ImageJ.

#### Serum traits measurement

The serum ALT and AST levels were measured according to the manufacturer’s instructions (Teco Diagnostics, NC9851324 for AST and NC9851323 for ALT tests). Serum total cholesterol levels and triglyceride levels were tested using Wako kits.

#### Single nuclei isolation from liver samples of FPC diet fed mice

For the snRNA-seq, we isolated the nuclei from male mice livers that fed with FPC diet for 16 weeks using a protocol modified from previous publications (8, 9). Briefly, frozen livers from 4 mice of liver-specific LBP KO group or 4 mice of WT group were cut and pooled according to the group. Samples were minced and transferred to the Dounce homogenizer. We added 4 ml of lysis buffer consisting of 10 mM Tris-HCl, 10 mM NaCl, 1 mM MgCl_2_, 0.1% IGEPAL and 0.3 U/μL RNase inhibitor to 80 mg pooled liver samples, then homogenized the tissue carefully. After filtering through 30 μm cell strainer, 4 ml of WRB buffer (PBS, 2% BSA, and 0.2 U/μL RNase inhibitor) was added and centrifuged at 500 g for 5 min at 4 °C. The lipid layer was removed, the pellet was resuspended in the WRB buffer and filtered again with 40 μm Flowmi cell strainer. Nuclei were pelleted by centrifuging at 500 g for 5 min at 4 °C again and resuspended in WRB buffer. To assess the quality of the isolated nuclei, we stained the nuclei with Hoechst stain for evaluation and counting.

#### 10x Genomics single-nucleus RNA sequencing

We performed 10x Genomics single-nucleus RNA sequencing at UCLA Technology Center for Genomics & Bioinformatics Core immediately after the nuclei isolation. Single cell RNA sequencing was prepared using 10X Genomics’ Chromium Next GEM Single Cell 3’ Kit v3.1 following the manufacturer’s protocol. The 10X 3’GEX libraries were sequenced on the Illumina NovaSeq6000 platform at a read length of 28x 90 and sequencing depth of 300-400 million reads per sample.

#### Single-nucleus RNA-sequencing analysis

Sequencing data de-multiplexing was performed using CellRanger mkfastq software v7.2.0 (10X Genomics). Sequenced reads in FASTQ format were aligned and analyzed via the CellRanger count pipeline v7.2.0 (10X Genomics) using the refdata-gex-mm10-2020-A mouse (10X Genomics) reference database under default settings. The R package Seurat was used to cluster the cells in the merged matrix (10). Subclusters were annotated using the marker genes according to previous publications (11–13).

#### Statistical analysis

Non-paired Student’s *t*-test or ANOVA were used to determine the statistical significance defined at *P* < 0.05. Data was shown as mean, and the error bar indicates SEM. Group sizes were based on statistical analysis of variance and prior experience with similar *in vivo* studies. \**P*<0.05; ** *P*<0.01; *** *P*<0.001; **** *P*<0.0001.

#### Study approval

All mouse experiments in this study were approved by the UCLA Institutional Animal Care and Research Advisory Committee, and performed in accordance with the recommendations in the Guide for the Care and Use of Laboratory Animals of the National Institutes of Health.

## Data availability

The sequencing data generated in this study can be accessed under GSE254626. The publicly available datasets we used were from the Gene Expression Omnibus at the National Center for Biotechnology Information (NCBI) or the Genotype-Tissue Expression (GTEx) project. The MASH patient dataset with disease grade is under GEO accession number GSE130970. The genome-wide association study is available from a study that measured blood proteins (14). The serum protein levels are available in the study of proteo-transcriptomic map of non-alcoholic fatty liver disease (15). The mouse liver single-cell RNA-seq data (*Tabula Muris*) (16) are available from the Gene Expression Omnibus (GSE109774). The human liver cellular landscape by single-cell RNA-seq is from GSE115469 (13).

## Acknowledgments

We thank all members of the Sallam laboratory and UCLA Lipidomics Core. The confocal microscopy was performed at the California NanoSystems Institute of Advanced Light Microscopy/Spectroscopy Facility at UCLA. We would like to thank Marcus Alvarez and Paivi Pajukanta for providing technical support of the single-nuclei isolation. T.S. is supported by National Institute of Health (NIH) grants (DK118086, HL139549, HL149766), an American Heart Association Transformational Project grant and Burroughs Wellcome Fund Career Award for Medical Scientists. D.W. is supported by American Heart Association Postdoctoral Fellowship (906049). A.G. is supported by R01 MD015904, K23 DK106528, R03 DK121025, ULTR001881/DK041301 (UCLA CURE/CTSI Pilot and Feasibility Study). Scientific schematics were generated with BioRender.

---

**Supplemental Figure 1.**
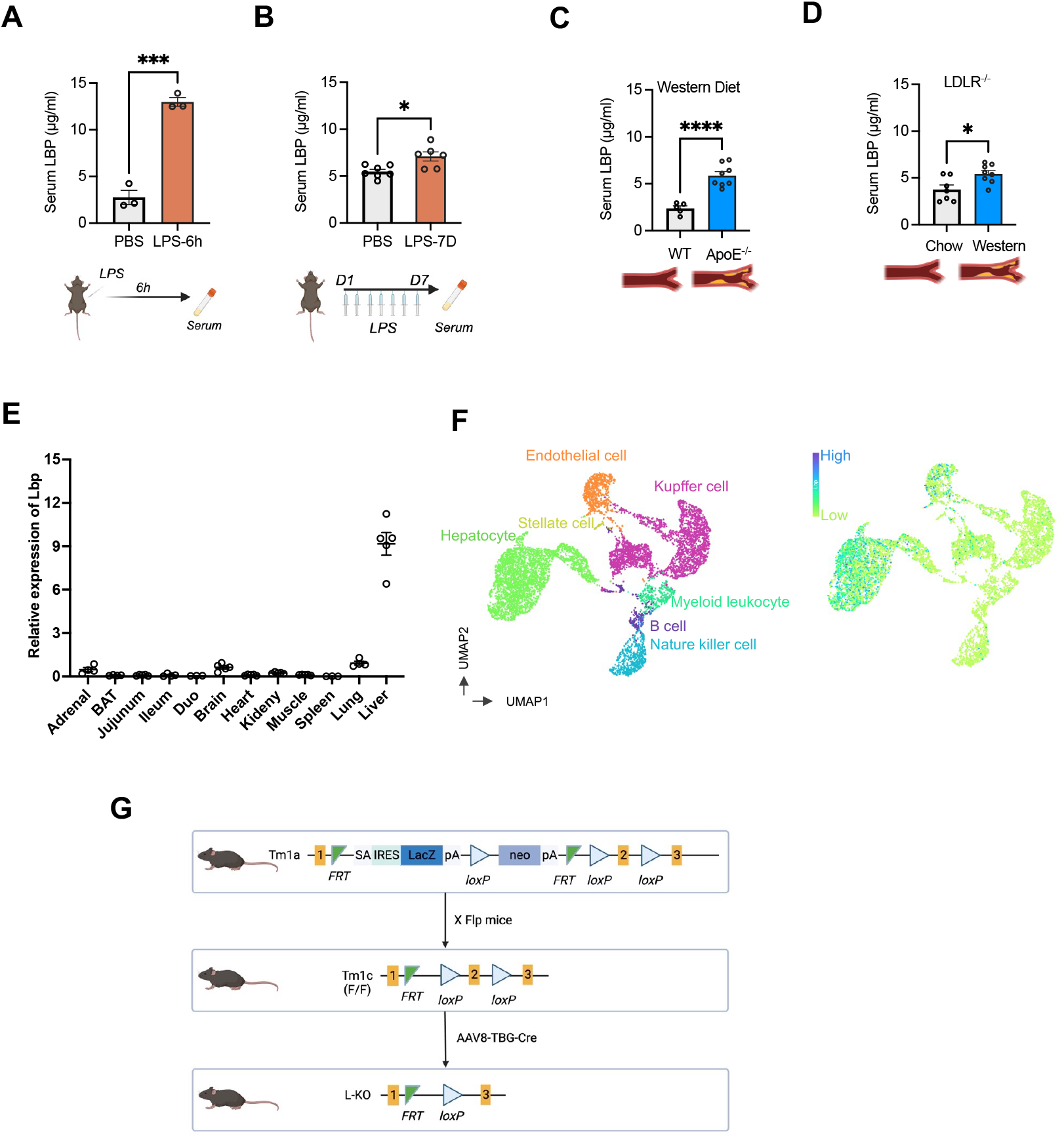
Circulating LBP is regulated with lipid-rich feeding. (A). Serum LBP 6 hours after LPS treatment (n= 3 per group). (B). Serum LBP after 1 week of LPS (n= 6∼7 per group). (C). Serum LBP from ApoE KO mice compared with WT on Western diet for 4-6 weeks (WT: n= 5, ApoE^-/-^: n= 8). (D). Serum LBP from LDLR KO mice fed with the Western diet for 16 weeks compared with chow diet (n= 7∼8 per group). (E). Gene expression of *Lbp* in a mouse tissue panel (n= 3∼5 per group). (F). Mouse liver single-cell RNA-seq data. UMAP was produced by cellxgene using GSE109774 dataset. (G). Strategy for generating LBP ^flox/flox^ mice. Data represent mean ± SEM. Non-paired Student’s *t*-test was applied. *: p < 0.05; ***: p < 0.001.

**Supplemental Figure 2.**
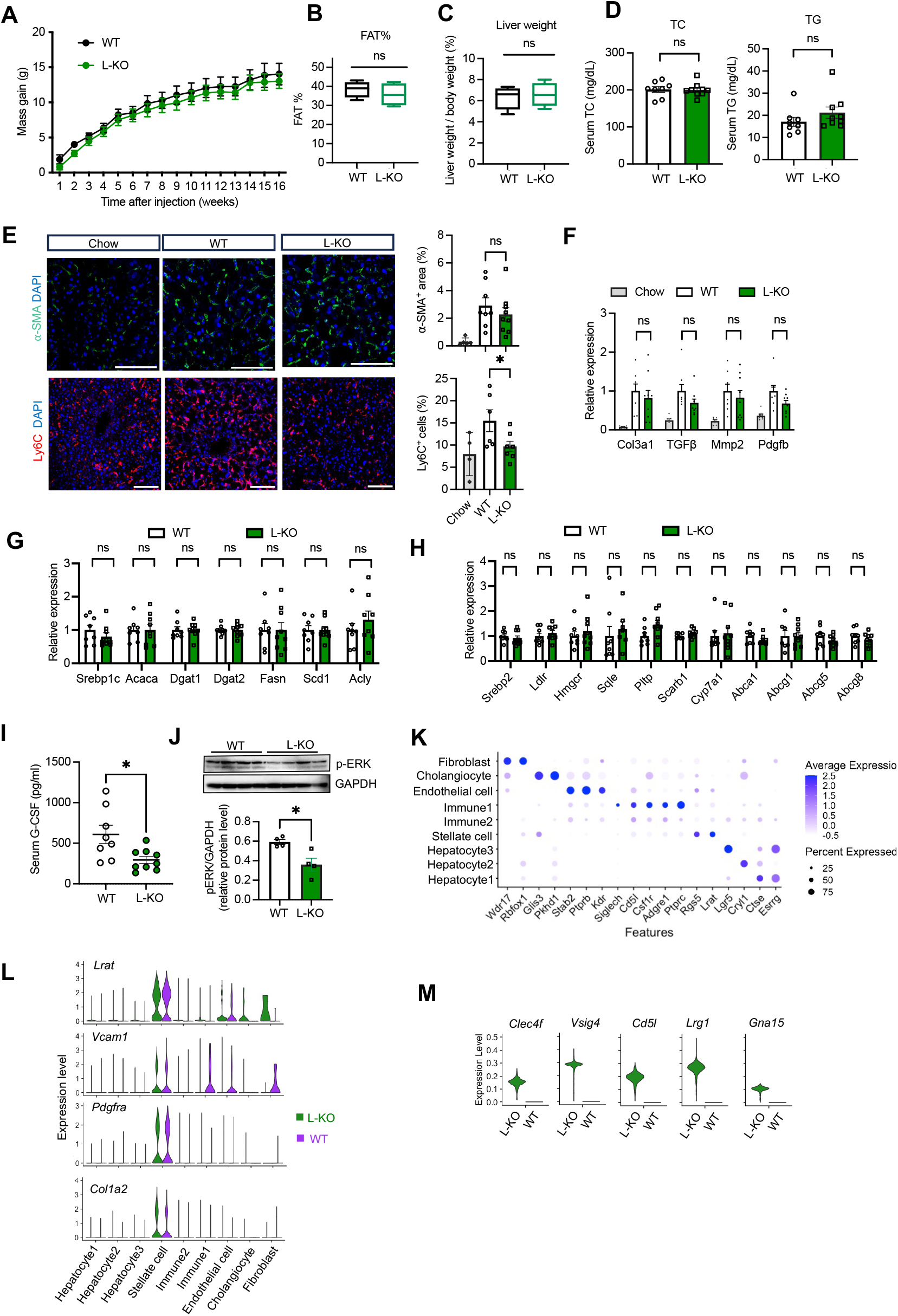
Mice lacking hepatocyte LBP exhibit reduced liver inflammation but not fibrosis. (A). Body weight comparing WT and L-KO mice on the FPC diet (n= 8∼9 per group). (B). Fat mass between WT and L-KO (n= 5 per group). (C). Liver/body weight ratios from B (n= 5 per group). (D). Serum lipid including total cholesterol (TC) and total triglyceride (TG) (n= 8∼9 per group). (E). αSMA staining and Ly6C staining from liver section (n= 8∼9 per group). Bar, 100 μm. (F). Gene expression of fibrosis markers (n= 8∼9 per group). (G). Gene expression measurements of triglycerides related pathways from mouse livers in WT and L-KO groups (n= 8∼9 per group). (H). Gene expression measurements of cholesterol related pathways from mouse livers in WT and L-KO groups (n= 8∼9 per group). (I). Serum G-CSF (n= 8∼9 per group). (J). Western blot of Phosphorated ERK (p-ERK) from whole liver lysates (*n*=4 per group). The bar plot shows the quantification results using ImageJ. (K). The expressions of marker genes from different clusters in snRNA-seq analysis in Figure 1. (L). Normalized expression of *Lrat, Vcam1, Pdgfra* and *Col1a2* in stellate cells under WT and L-KO group in snRNA-seq data. (M). Violin plots showing the expression of representative genes in WT and L-KO mice. Data represent mean ± SEM. Non-paired Student’s *t*-test was applied. *: p < 0.05; ns: not significant.

**Supplemental Figure 3.**
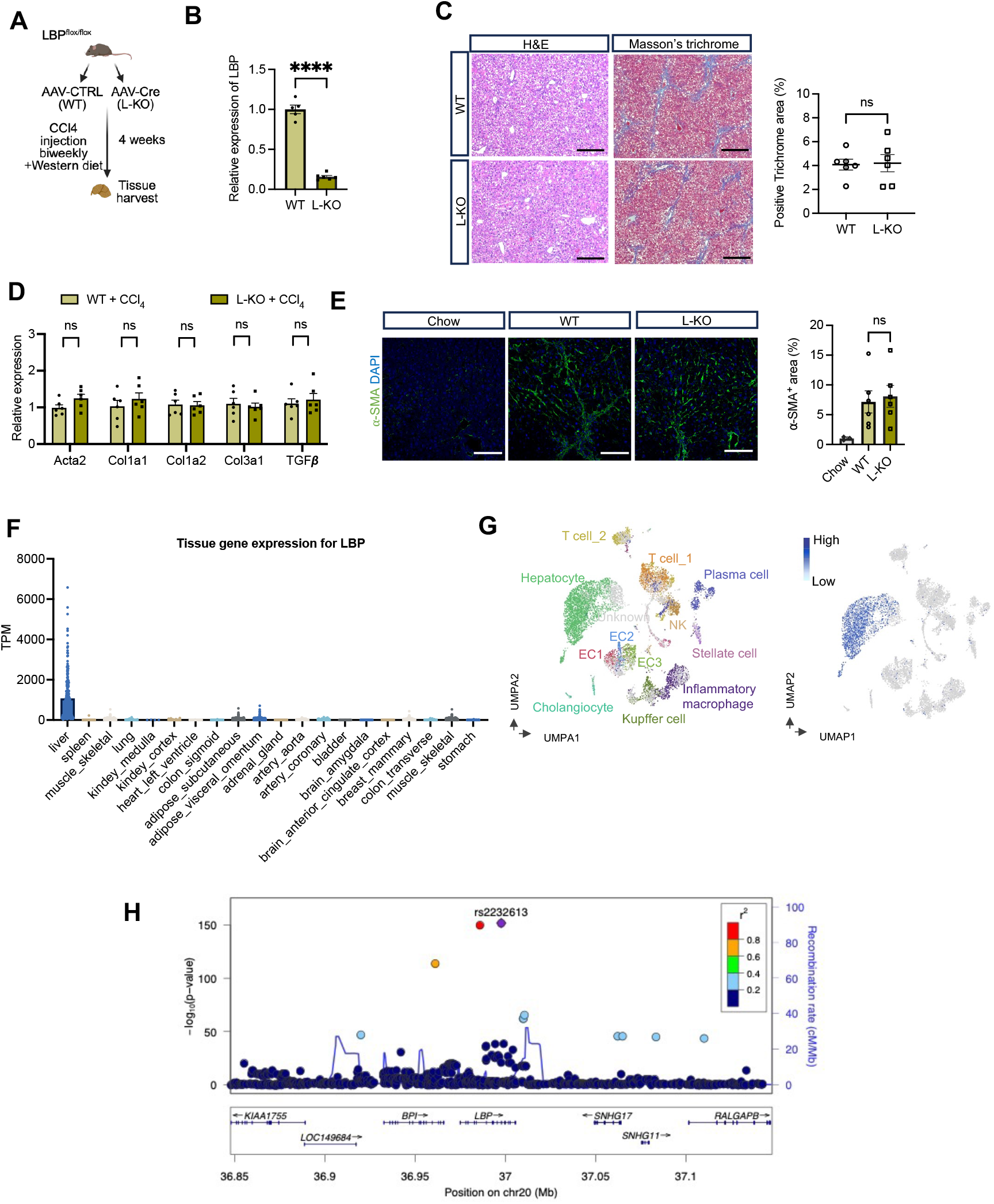
Variants at human LBP are associated with inflammation. (A). Experimental design of CCl_4_ treatment. (B). RT-qPCR results from liver in A. (n= 6 per group). (C). H&E staining and Masson’s trichrome staining from liver sections. Quantification of Masson’s trichrome staining area shown (n= 6 per group). Bar, 100 μm. (D). Gene expression of fibrosis markers in liver (n= 6 per group). (E). Staining of αSMA in liver section with quantification (n= 4 in chow, and n= 6 per group in treatment). Bar, 100 μm. (F). LBP expression in human tissues from GETx database. (G). Human liver single-cell RNA-seq. UMAP produced by Single Cell Expression Atlas using GSE115469 dataset. (H). LocusZoom plot showing genome-wide significant association observed in serum IL15 levels at LBP. Data represent mean ± SEM. Non-paired Student’s *t*-test was applied. ****: p < 0.0001; ns: not significant.

**Supplemental Table 1:**
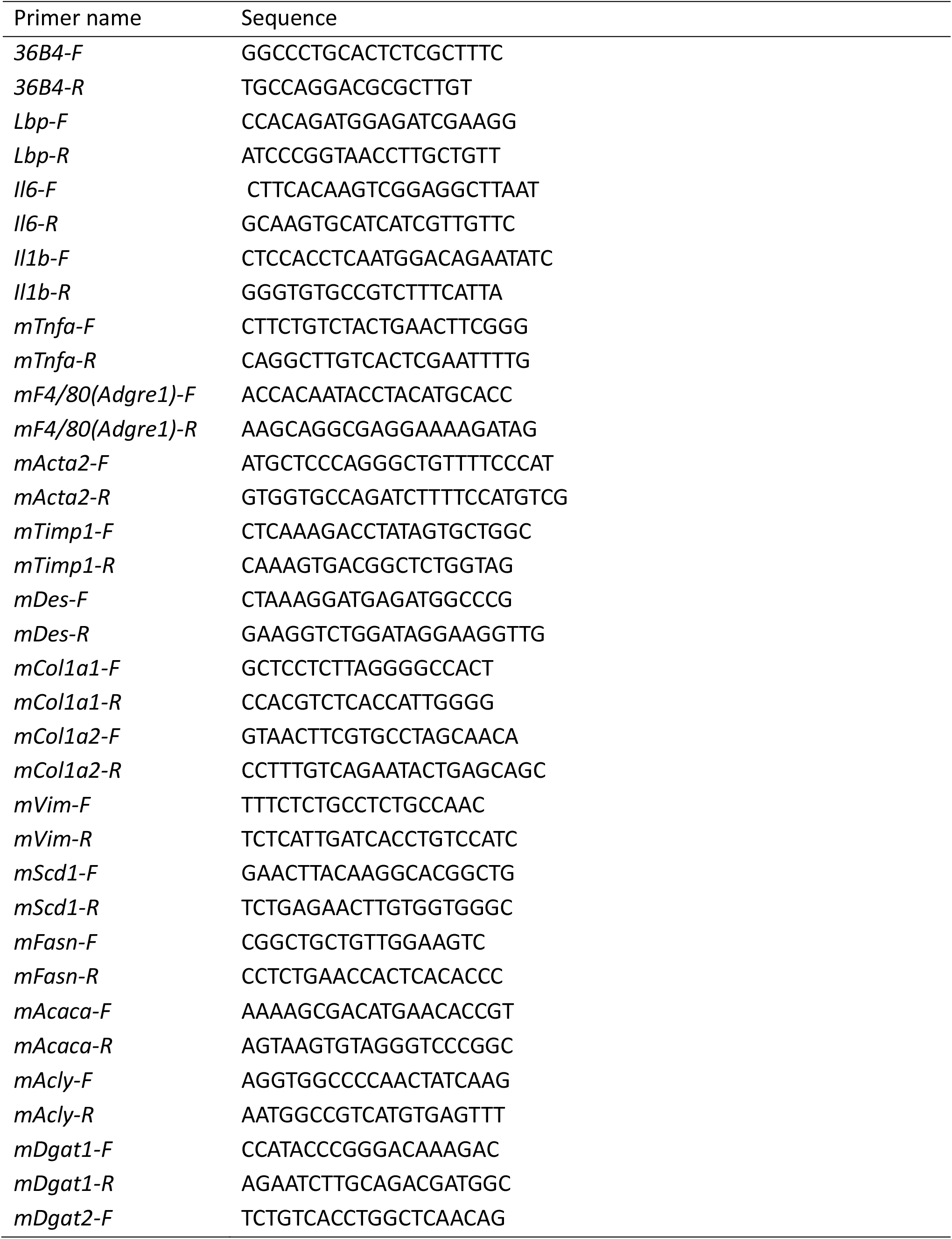
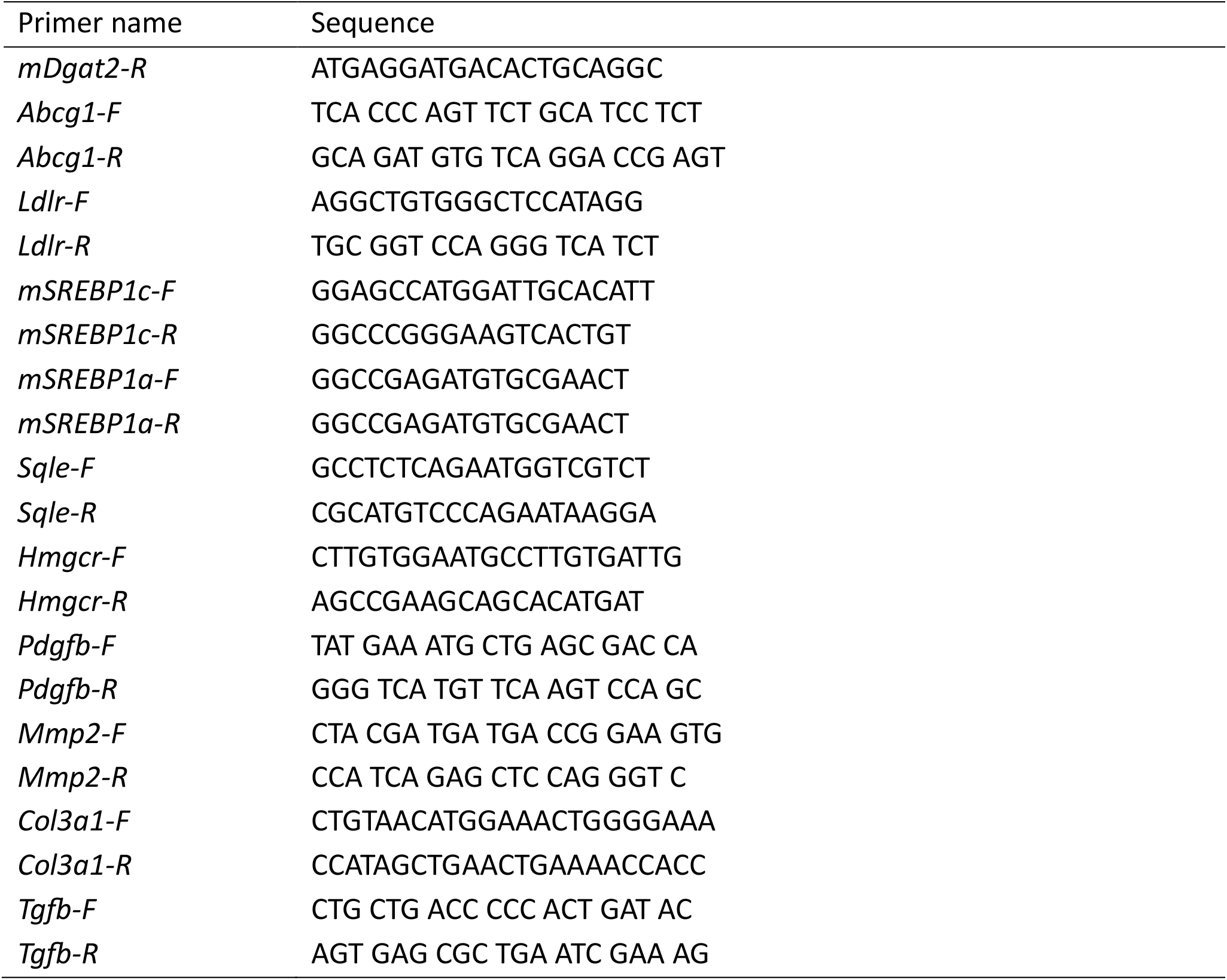
RT-qPCR primers.

